# MiRNA Atlas: A Literature-Derived Database of MicroRNAs Bridging Osteoarthritis and Appendage Regeneration

**DOI:** 10.64898/2026.09.17.748327

**Authors:** Abigail T. Parker, Virginia Byers Kraus

## Abstract

1.

Many microRNAs (miRNAs) regulate tissue remodeling, cellular plasticity, and repair across evolutionarily distant vertebrate lineages that are capable of regenerating appendages such as amputated limbs, fins, and antlers, as well as in human articular cartilage responding to injury. These miRNAs often belong to the same families and exert conserved, though occasionally inverted, regulatory effects. This strong cross-species overlap motivates the present literature-derived analysis. Osteoarthritis (OA), the most prevalent joint disease, still lacks disease-modifying therapies, in part because of the longstanding assumption that adult mammalian cartilage demonstrates no intrinsic reparative capacity. Yet human cartilage retains a latent repair program activated by mechanical and inflammatory stress. Some injured or degenerating joints may never be clinically recognized as osteoarthritic because their intrinsic repair capacity is sufficient to restore tissue integrity; in others, where repair capacity is diminished or damage exceeds it, the repair program is insufficient and OA becomes clinically manifest. MiRNAs are established post-transcriptional regulators of cartilage homeostasis, degeneration, and appendage regeneration, yet because the OA and regeneration research fields have advanced largely independently, the insights available at their intersection have gone unrecognized. To close this gap, we systematically mined both literatures to construct an auto-updating, cross-referenced atlas of OA- and appendage regeneration-associated miRNAs. Integrating these datasets identified a core set of shared miRNA families, delineated miRNAs unique to each field, and mapped convergent families onto common pathways governing matrix remodeling, dedifferentiation, senescence, and inflammation. We propose that regeneration-competent species can inform the identification of therapeutic miRNAs, such as miR-133, miR-21, and let-7, capable of activating endogenous cartilage repair. Collectively, this synthesis and its accompanying web-based miRNA atlas (https://mirnaatlas.shinyapps.io/mirnaatlas/) establish a comparative framework for regenerative miRNA biology and provide a continually updated resource to accelerate discovery of disease-modifying, RNA-based therapies for OA.

**Graphical Abstract:** **Figure 1.**
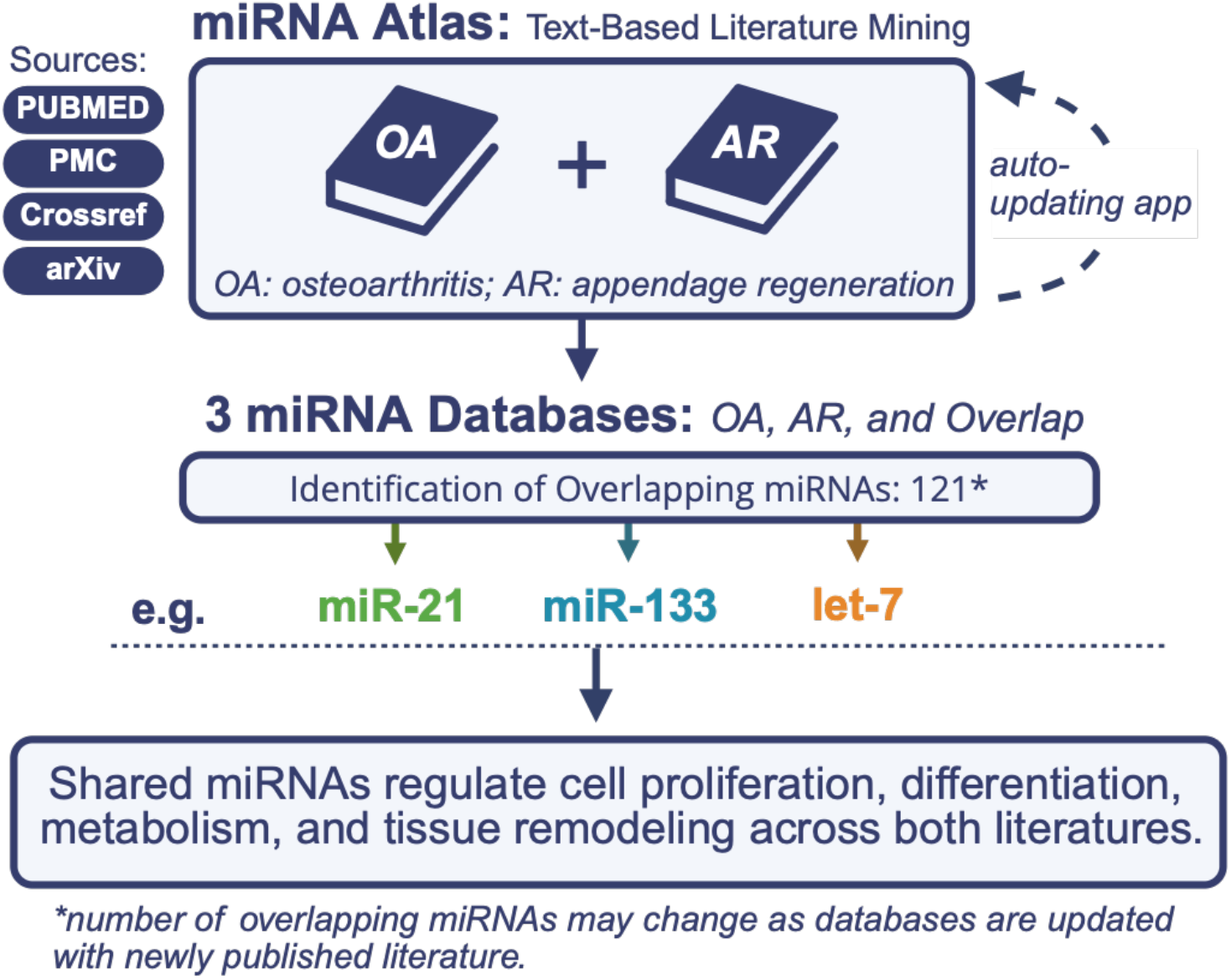
Graphical Abstract. The miRNA Atlas is a web-based, continually updated literature-mining application that indexes PubMed, PubMed Central, Crossref, and arXiv to identify shared microRNA regulators in studies of osteoarthritis and appendage regeneration. We highlight three candidates involved in both osteoarthritis (OA) and appendage regeneration (AR) with therapeutic potential to enhance innate regenerative capacity.

## 3. Introduction

### 3.1 Conserved regeneration, divergent capacity

Responses to injury can vary widely across tissues. For instance, some tissues can fully regenerate, restoring the original structure, while others heal with residual scarring and tissue dysfunction. Mechanistically, both reparative outcomes involve similar early responses, including injury sensing, inflammation, cell proliferation, extracellular matrix remodeling, and the reactivation of developmental programs. Successful regeneration, however, requires the sustained coordination of these responses, enabling progenitor cells to expand and differentiate while reestablishing tissue organization and function.

These differences in regenerative capabilities are particularly evident across vertebrates, where the same injury can yield dramatically different outcomes depending on intrinsic regenerative competence. Urodele amphibians, such as the axolotl (*Ambystoma mexicanum*), regenerate fully patterned limbs containing skin, muscle, bone, cartilage, and nerves, through a proliferative blastema of transiently dedifferentiated cells (1,2). Zebrafish execute the same regenerative program on a faster timescale, regenerating amputated fins within days through a blastema-mediated process that restores both bony rays and the surrounding soft tissue (3). Conversely, mammalian joints lie toward the opposite end of this regenerative spectrum, with little capacity for complete restoration after injury.

However, even in mammals, the prevailing view––that regeneration is largely restricted or–– underestimates their true capacity. Deer antlers regenerate repeatedly and completely, with annual regrowth driven by stem cells through a developmental program that recapitulates endochondral ossification and cartilage formation (4). Humans can regenerate substantial portions of the liver following partial hepatectomy (5,6) and can restore the distal tips of amputated digits under favorable conditions (7,8). The MRL/MpJ mouse, initially identified for its ability to heal ear-punch wounds without fibrosis, also regenerates distal digit tips and demonstrates an enhanced articular cartilage repair response following joint injury compared with conventional inbred strains (9–11). Collectively, these examples demonstrate that regenerative capacity is not binary but exists along a continuum, reflecting the degree to which evolutionarily conserved repair programs are activated across species, tissues, and anatomical contexts. Given that these organisms diverged evolutionarily hundreds of millions of years ago, pathways shared between highly regenerative species and mammals likely represent ancient mechanisms predating modern vertebrate lineages. Comparative analyses across diverse systems, therefore, provide a powerful approach for distinguishing core regulators of regeneration from species-specific adaptations.

### 3.2. Appendage regeneration as a model of coordinated, whole-organ repair

Successful regeneration and incomplete repair are distinguished less by which processes occur than by whether each unfolds in the correct spatial and temporal contexts (12). Their coordination enables injured tissue to progress from an initial repair response to full restoration of structure and function, suggesting that successful regeneration depends on regulatory mechanisms that integrate multiple cell types and biological processes rather than acting through isolated pathways. Appendage regeneration is a particularly powerful model of this coordination. Regrowing an appendage requires rebuilding a spatially patterned organ, including cartilage, bone, muscle, connective tissue, blood vessels, nerves, and epithelium (13,14). Instead of only replacing individual lost cells, the organism must reestablish tissue architecture and function across the entire appendage. Osteoarthritis (OA) poses a related challenge, affecting the complex, multi-tissue environment of the joint: cartilage, subchondral bone, synovium, and surrounding soft tissues. Similar to a regenerating limb, the osteoarthritic joint triggers endogenous repair mechanisms. However, because the joint’s repair ability is limited or because the damage is too extensive, these responses often do not fully restore the original tissue structure and function.

Articular cartilage has traditionally been regarded as a tissue with minimal capacity for self-repair. This assumption is among the oldest in orthopedics, dating to at least William Hunter’s 1743 observation that “cartilage injury is a troublesome thing and, once injured, is seldom repaired” (15). The hypocellular, avascular, and aneural characteristics of articular cartilage have reinforced a prevailing bias in both clinical and basic science literature that cartilage lacks meaningful regenerative capacity (16). Clinically, this perspective is evident in existing repair strategies such as microfracture and autologous chondrocyte implantation, which aim to augment or replace damaged tissue rather than restore or enhance its endogenous self-repair capacity. However, the inability to fully regenerate should not be equated with the absence of endogenous repair mechanisms. Instead, human articular cartilage mounts a coordinated repair response to injury and osteoarthritic damage, including matrix synthesis, cellular proliferation, progenitor-cell activation, and phenotypic adaptation.

Where this response keeps pace with damage, degeneration may never progress to clinically recognizable disease; the OA we diagnose reflects those joints in which repair is outpaced or insufficient.

The evidence for this endogenous response is first apparent morphologically. Sites of cartilage degeneration are characteristically bordered by chondrocyte clusters in which resident cells escape their normal single-lacunar arrangement and proliferate together; these clusters display molecular markers of progenitor cells rather than terminally differentiated chondrocytes (17). Progenitor-like cells are also mobilized more broadly at lesion edges, migrating into and around damaged regions (18) in a pattern that mirrors the wound-proximal progenitor recruitment observed during appendage regeneration (19,20), though at markedly lower efficiency. Thus, although OA cartilage does not form a blastema or regrow a patterned appendage, injury still elicits local cellular behaviors consistent with an attempt to restore the damaged tissue.

Single-cell profiling of human OA cartilage shows that this reparative response is reflected in distinct chondrocyte states. Chou et al. (2020) identified seven chondrocyte phenotypes across intact and damaged regions of OA cartilage (21). Non-diseased tissue was dominated by homeostatic and hypertrophic chondrocytes, whereas diseased regions were enriched for regulatory, reparative, pre-fibrochondrocyte, and fibrochondrocyte populations. These populations showed distinct functional signatures: reparative chondrocytes expressed *COL2A1, CILP, COL3A1*, and *COMP*; pre-fibrochondrocytes expressed *IL11, COL2A1, CILP*, and *OGN*; fibrochondrocytes expressed *COL1A1, COL1A2, S100A4*, and *PRG4*, consistent with extracellular matrix organization; and regulatory chondrocytes expressed *CHI3L1* and *CHI3L2*. Consistent with these findings, Ji et al. (2019) identified cartilage progenitor cells and proliferative chondrocytes in human OA cartilage, alongside regulatory and fibrocartilage chondrocyte populations (22); the chondrocyte progenitor population expressed discrete cell state markers (*BIRC5, CENPU, UBE2C, DHFR* and *STMN1)* and was proposed to contribute to cartilage maintenance and repair, further supporting the presence of reparative cells in human OA cartilage. Collectively, these findings indicate that OA cartilage retains multiple components of an endogenous reparative program, including progenitor activation, anabolic matrix synthesis, and the emergence of reparative chondrocyte states. However, whether a joint progresses to OA may depend on the balance between these reparative responses and the persistent inflammatory, catabolic, and fibrogenic programs with which they coexist; when repair keeps pace, degeneration may never reach clinical recognition, whereas OA marks the trajectory in which repair is engaged but ultimately overcome, resulting in a failure to restore native cartilage architecture. Consequently, OA may represent an incompletely activated or poorly sustained regenerative state rather than a complete absence of regeneration, suggesting a therapeutic strategy focused on reinforcing endogenous repair and tipping this balance toward true regeneration.

### 3.3. MicroRNAs are master regulators of tissue maintenance and regeneration

MicroRNAs (miRNAs, or miRs) are a class of small, non-coding RNAs that post-transcriptionally regulate gene expression by guiding the RNA-induced silencing complex (RISC) to complementary sequences, typically in the 3′ untranslated regions of target mRNAs (23,24). MiRNAs are also among the most evolutionarily conserved regulatory elements in the genome (25,26). Because miRNA activity depends on sequence complementarity, a conserved miRNA sequence can preserve recognition of conserved target transcripts and regulatory pathways across species. A single miRNA can target dozens to hundreds of transcripts, allowing miRNAs to regulate entire gene networks rather than simply individual genes. Accordingly, miRNA-mediated regulation is well suited to the large-scale, multi-gene transitions underlying cell fate change, dedifferentiation, tissue remodeling, and regeneration.

In healthy cartilage, miRNAs maintain chondrocyte identity and extracellular matrix homeostasis; in OA, disruption of these networks shifts the balance from anabolic to catabolic programs and drives tissue degradation (27–29). Beyond maintenance, miRNAs also facilitate an active repair response to osteoarthritic injury. Hsueh, Önnerfjord, and Kraus (2025) identified a program of 69 distinct small RNAs (smRNA) induced under osteoarthritic stress that drives cartilage matrix protein anabolism and anabolic gene expression (30). This program follows an anatomical gradient, which is strongest in ankle OA cartilage, intermediate in knee OA cartilage, and weakest in hip OA cartilage. This distal-to-proximal pattern mirrors the greater regenerative competence of distal structures in regenerative organisms, raising the possibility that anatomical variation in regenerative capacity is itself shaped by smRNA regulation. Several of these smRNAs are strongly conserved, at both the sequence and expression levels, with regenerative programs identified in axolotl, bichir, and zebrafish (31,32).

Notably, smRNAs encompass not only miRNAs but also PIWI-interacting RNAs (piRNAs) and other non-coding RNA species, and the regenerative signal is unlikely to be confined to miRNAs alone. We have identified regenerative piRNAs in human cartilage, indicating that multiple smRNA classes participate in the endogenous repair response. The present literature-mining workflow focuses on miRNAs because, to date, they are the only smRNA class sequenced in limb-regenerating organisms; extending this comparative work to piRNAs and other smRNAs will be an important direction for future study. These observations indicate a shared smRNA regulatory signature spanning cartilage repair and appendage regeneration. Despite this overlap, the OA and regeneration literatures have developed separately, even though both focus on miRNA regulation of cell phenotype, proliferation, differentiation, and tissue remodeling. OA studies typically frame these mechanisms in the context of cartilage homeostasis and disease, whereas regeneration studies emphasize blastema formation, tissue patterning, and restoration of tissue integrity. The degree of overlap between these conserved miRNA programs in OA and regeneration has not been systematically examined. Accordingly, we mined both literatures to identify shared miRNAs, construct a systematic atlas of their reported associations, and nominate candidates to therapeutically sustain cartilage repair.

## 4. Methods

### 4.1 miRNA Atlas: A Literature-Derived Integrated Database for Osteoarthritis- and Appendage Regeneration-Associated miRNAs

A literature-mining tool was developed to investigate the overlap of regulatory microRNAs between the appendage regeneration and OA literatures. The miRNA Atlas is a web application built in R using the Shiny framework that systematically gathers and cross-references miRNA publications, including articles and preprints, on OA pathophysiology and limb or appendage regeneration from four open-source databases. It extracts structured, machine-readable information from relevant publications and organizes the findings into three linked, searchable, and downloadable databases: an Osteoarthritis database, an Appendage Regeneration database, and an Overlap database. Rather than relying on a single manual literature search, the miRNA Atlas integrates published evidence across these distinct literatures, enabling comparison across studies that often differ widely in terminology, model systems, and experimental methods. The automated approach provides a reproducible system for identifying, organizing, and comparing miRNA findings, and it allows the underlying literature to be re-queried and updated over time.

### 4.2. Literature Mining and Database Construction

During each refresh cycle, the miRNA Atlas queries four free, publicly available literature application programming interfaces (APIs) in parallel: PubMed, PMC (which also indexes bioRxiv and medRxiv preprints), Crossref, and arXiv. We excluded proprietary or usage-metered databases such as Web of Science and Scopus to ensure that the tool remains freely reproducible and portable, without dependence on institutional licensing. Each source is queried using a Boolean combination of miRNA-related terms (e.g. microRNA, miRNA, miR, micro-RNA, and small non-coding RNA), and topic-specific terms. For OA, these include osteoarthritis, osteoarthrosis, degenerative joint disease, cartilage degeneration, and chondrocyte senescence; for appendage regeneration, these include limb, fin, tail, digit and digit-tip, and antler regeneration, as well as epimorphic regeneration and blastema. Query strings are intentionally left unquoted so that each API can apply its native term-mapping and synonym-expansion functions, substantially increasing recall relative to literal phrase matching. Because several terms central to appendage regeneration have unrelated meanings in adjacent fields, additional exclusion rules were applied to prevent topic misclassification. “Blastema,” which describes the mass of dedifferentiated cells during appendage regeneration, is also the name of a histological subtype of a pediatric nephroblastoma (Wilms tumor); citations discussing renal or nephroblastoma blastema were excluded from the regeneration database. Similarly, “appendage” was excluded when used in its dermatological sense (e.g., “skin appendage,” referring to hair follicles and sweat glands) rather than as a limb, fin, or tail structure, and “limb” was excluded from disease-name compounds such as limb-girdle muscular dystrophy that do not refer to an anatomical limb undergoing regrowth. Records returned by multiple sources are deduplicated using DOI or normalized title, retaining all contributing sources on the merged record and preserving the most complete available abstract.

Because free-text search engines such as Crossref rely on loose keyword co-occurrence, every candidate citation passes through a relevance gate before being added to a database. Using regular-expression matching, the title and abstract must show that the citation discusses both a miRNA and the topic of interest. For appendage regeneration, we add a further requirement that an anatomical structure, such as a limb, fin, digit, antler, blastema, or a recognized synonym, be explicitly named, to exclude the much larger body of literature related to generic tissue repair and wound healing. Mature and precursor miRNA identifiers are then identified with a regular expression that uses explicit boundary checks to prevent partial or spurious matches. This pattern recognizes conventional forms (miR-140, miR-140-5p, hsa-miR-140, let-7a, microRNA-21), as well as compressed list notations such as “miR-99a/100” and “miR-140, -146, -155.” Identifiers are normalized to a canonical form so that spelling and formatting variants of the same miRNA are treated as identical. Scored, dictionary-based classifiers annotate each citation for: species or model organism; joint or appendage studied; tissue or cell type; experimental model type (clinical, *in vivo, in vitro, ex vivo*, and computational); and experimental condition or injury paradigm, e.g., destabilization of the medial meniscus, monoiodoacetate injection, cytokine stimulation, or surgical amputation. The tool assigns regulation direction (upregulated, downregulated, or both) computationally from sentence-level language rather than from the underlying expression data. For example, splitting “miR-140 was downregulated, whereas miR-146a was increased” into clauses attributes the downregulation to miR-140 and the upregulation to miR-146a—an annotation that should be treated as provisional rather than verified. Because this classification relies on natural-language cues (e.g., “increased,” “suppressed,” “knockdown”), it can misattribute direction in cases of complex or conditional phrasing, multi-timepoint or multi-tissue findings, or negated statements. We recommend manually cross-checking any reported direction of regulation against the primary literature before using it to support a specific claim. For open-access citations, the pipeline additionally retrieves full-text XML and repeats miRNA extraction across the entire article, including tables and figure legends, where comprehensive lists of differentially expressed miRNAs are often reported. The Methods sections are also mined specifically to recover species, tissue, and model-type annotations that abstracts frequently omit.

The annotated citations are compiled into two long-format databases in which each row represents one miRNA–citation entry. A third, derived database is constructed by intersecting the two on miRNA identity, matched at the precursor level after stripping the mature-strand suffix and any precursor copy number. All three databases, along with the underlying raw citation metadata, are stored in a SQLite database and can be exported as CSV and Excel workbooks. The application refreshes automatically at set intervals to ensure recently published literature can continue to be indexed.

### 4.3. Utility of the miRNA atlas

The miRNA Atlas enables systematic, filterable, and reproducible comparisons of miRNA involvement across two fields of skeletal and regenerative biology that are rarely cross-referenced in the primary literature. The Overlap database identifies miRNAs reported in both the OA and appendage-regeneration literatures, making it directly useful for nominating candidate regulators shared between cartilage degeneration and pro-regenerative programs for limb, fin, and digit-tip regrowth. Each association is annotated with species, tissue, model type, and experimental condition or injury paradigm, enabling comparisons across biological and experimental contexts at scale. The rule-based extraction ensures that every entry is linked to a citation, maintaining a verifiable evidence base and avoiding hallucinated associations that can arise with unsupervised or generative summarization. Furthermore, because the tool re-queries the literature on an ongoing basis rather than relying on a single point-in-time search, its evidence base remains current as new research emerges in both fields.

## 5. Research Applications: Mining the miRNA Literature for Shared Regulators of Degeneration and Regeneration

### 5.1 A Comparative Atlas Reveals a Shared miRNA Signature

To characterize the miRNA literatures underlying OA and appendage regeneration, we compiled a comparative atlas capturing publication trends, unique miRNAs, and shared regulatory miRNAs across both fields. To date, this atlas has identified 1,610 miRNA-associated citations with 1,622 unique OA miRNAs drawn from 8,920 database entries; the atlas also identified 40 appendage regeneration citations with 160 unique miRNAs drawn from 247 database entries. The overlap between the two conditions includes 1233 citations and 121 unique miRNAs. Annual publication trends reveal the substantial growth in OA-related miRNA literature since 2008, with citations rising sharply after 2014 and peaking in recent years, while appendage regeneration research has remained comparatively smaller and stable throughout the same period (Fig. 1A). Despite this imbalance in publication volume, both fields converge on a shared set of miRNA regulators. Ranking the top unique miRNAs by citation count highlights field-specific candidates as well as strongly overlapping miRNA candidates such as let-7, miR-21, and miR-133. These overlapping miRNA regulators represent high-priority targets for future mechanistic studies, as they may point to conserved regulatory pathways governing tissue degeneration and regenerative capacity alike.

**Figure 2.**
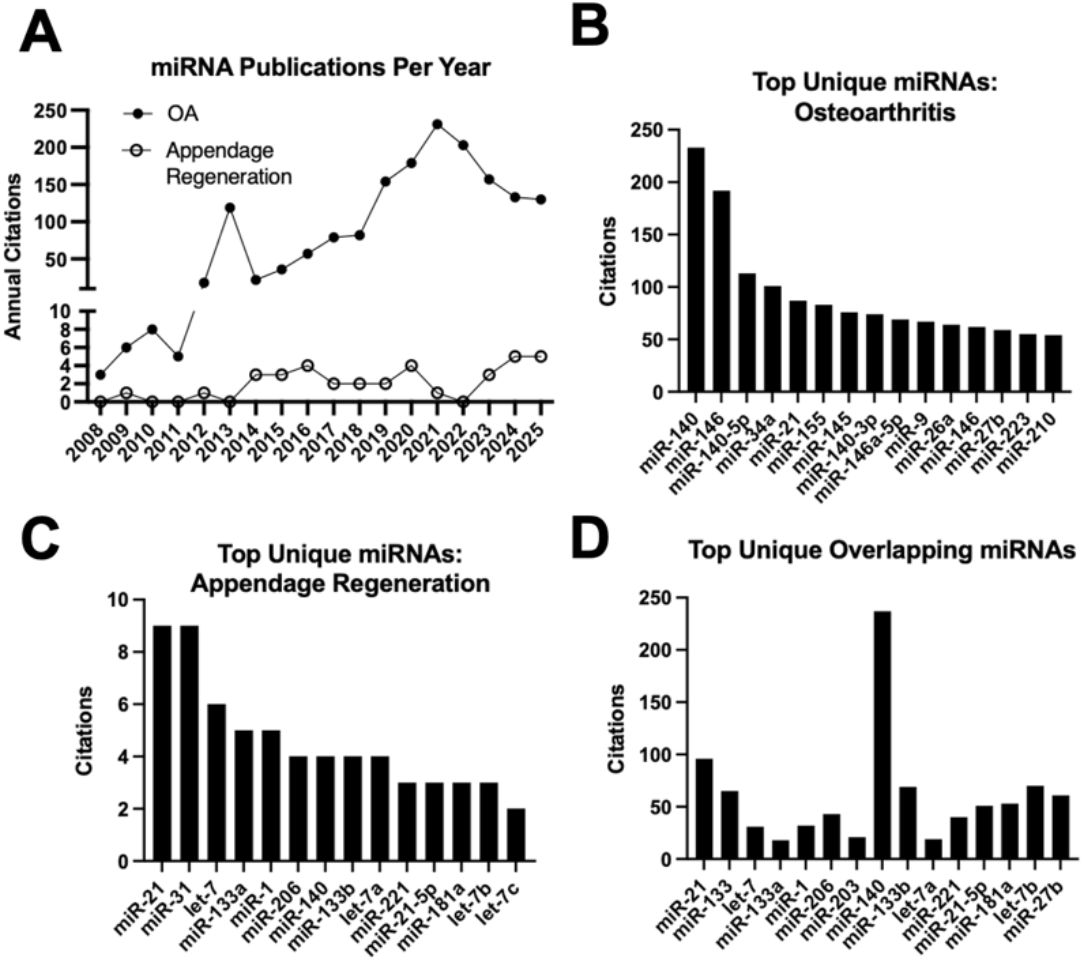
Comparative analysis of miRNA research trends and unique and overlapping miRNAs in osteoarthritis (OA) and appendage regeneration literatures using the miRNA Atlas. **(A)** Annual number of miRNA-related publications (citations) from 2008–2025 for OA (closed circles) and appendage regeneration (open circles), illustrating the respective growth in research output between the two fields. **(B)** Citation counts for the top unique miRNAs associated with osteoarthritis, ranked from highest to lowest. **(C)** Citation counts for the top unique miRNAs associated with appendage regeneration, ranked from highest to lowest. **(D)** Citation counts for the top miRNAs found to overlap between both OA and appendage regeneration literature, ranked by the degree of overlap, highlighting candidate miRNAs with potential shared regulatory roles across both processes.

Beyond ranking, the atlas allows each shared regulator to be examined individually, tracing how a single miRNA has been studied across both degenerative and regenerative contexts. Rather than treating overlap as a summary statistic, it permits further investigation into why a given miRNA recurs in both literatures and what conserved role that recurrence implies for cartilage. The candidates below were independently prioritized in the OA and appendage-regeneration literature, but the two fields have approached them from opposite directions. OA studies have focused on how their dysregulation contributes to cartilage breakdown, while regeneration studies have asked how their activity supports tissue rebuilding. Considered together, these two related research areas suggest a clear, testable model of miRNA function that is not evident when examining each field separately.

### 5.2. miR-21

In vertebrate species whose evolutionary lineages diverged over 420 million years ago, miR- 21 is a consistent regulatory miRNA involved in the regenerative response. In blastemas from the zebrafish caudal fin, bichir pectoral fin, and axolotl forelimb, miR-21 is the most abundantly expressed miRNA across all three species (32). As miR-21 expression increases during limb regeneration, it targets *tgfbr2*, an important mediator of TGFβ signaling that regulates cell proliferation, differentiation, extracellular matrix remodeling, and tissue repair (32). miR-21 has also been shown to target *jagged1* during axolotl limb regeneration. Jagged1 is a Notch ligand involved in progenitor cell maintenance, and its repression during axolotl limb regeneration may aid the transition from proliferative blastemal cells toward differentiation (33).

In osteoarthritis, miR-21 plays a more context-dependent role, and studies have not resolved whether it acts as an insufficient reparative signal or as an active driver of OA pathogenesis. Evidence for a deleterious role comes largely from settings of sustained elevation in stressed chondrocytes and inflamed joint tissue. There, miR-21 represses GDF-5 (34) and Spry1 (35), promoting cartilage matrix degradation and angiogenesis. Furthermore, its deletion attenuates cartilage breakdown in models of temporomandibular joint OA (35). Beyond cartilage, synovial fibroblasts package miR-21-5p into extracellular vesicles that carry inflammatory signals to chondrocytes (36), and some extracellular miR-21 has been shown to sustain OA-associated pain as a TLR7 ligand (37). Other work has also described protective roles of miR-21, including promoting hyaline cartilage production in IL-1β-stressed chondrocytes (38), alleviating OA by inhibiting inflammatory factors and altering macrophage phenotype (39), and appearing among the beneficial cargo contents of extracellular vesicles released by mesenchymal stromal cells (40). Hsueh and colleagues identified miR-21 within a 69-smRNA program induced by osteoarthritic stress that promotes matrix anabolism, reduces matrix degradation, and suppresses inflammatory cytokine secretion as described above. Induction of miR-21 overexpression by viral vector delivery to cultured chondrocytes subsequently increased the expression of cartilage anabolic factors. Notably, miR-21 expression in osteoarthritic cartilage samples exhibits an anatomical gradient (ankle > knee > hip), resembling the patterns seen in three regenerating appendage species. This links the human cartilage miRNA program to the conserved regenerative responses observed across vertebrates (30). These outcomes emphasize the importance of the cellular context in which miR-21 is expressed, the targets available to it, and the duration of its elevation. Chronic accumulation in stressed chondrocytes or inflamed synovium appears pathological, whereas transient elevation within a coordinated reparative program, or delivery from progenitor and resolving-macrophage populations, appears beneficial.

### 5.3. miR-133

miR-133 has repeatedly emerged as a shared regulator of cell state and proliferation in appendage regeneration and OA. In regenerative tissues, miR-133 expression is reduced as cells enter a proliferative state; similarly, in OA studies, miR-133 is associated with altered chondrocyte proliferation, survival, and differentiation. As the regeneration literature spans multiple vertebrate species, cross-species comparisons allow investigation of both sequence homology and the extent to which miR-133 function is conserved or species-specific. Mature miR-133a differs by no more than a terminal nucleotide among zebrafish, newt, lizard, mouse, and human, with completely conserved seed sequences; therefore, functional differences across these systems are more likely to arise from cellular context, expression level, and target availability.

Studies across regenerative vertebrates have examined miR-133 during the early stages of regeneration, with several emphasizing its depletion in temporal association with blastemal proliferation. Following zebrafish caudal fin amputation, Fgf signaling from the wound epidermis rapidly reduces miR-133; in contrast, maintaining miR-133 suppresses blastemal proliferation and stalls regenerative outgrowth (41). In newt tail and caudal spinal cord regeneration, miR-133a decreases in ependymal cells surrounding the central canal, relieving repression of RARβ2 and permitting retinoid signaling (42). miR-1 and miR-133a are encoded in the same cluster and target the RARβ2 3′-UTR, while RARβ2 suppresses both miRNAs. This negative feedback loop sustains RARβ2 signaling during ependymal tube outgrowth (43). Integrated microRNAome, transcriptome, and proteome profiling in newts (*Cynops orientalis*) similarly identifies miR-133a among the miRNAs downregulated during blastema formation and early redevelopment, which describes the recurrence of events and processes necessary for embryonic development (44). As miR-133a decreases, repression of its target *G6PD* is relieved, increasing *G6PD* expression and supporting pentose phosphate pathway (PPP) flux and the metabolic and nucleotide synthesis demands of proliferating blastemal cells (45). In *Anolis carolinensis*, miR-133a and miR- 133b are elevated in the regenerating tail base relative to the tail tip (46). Consistent with these differential cellular conditions, the tail tip contains the hypo-differentiated proliferative growth zone, whereas the tail base shows a more advanced state of cell differentiation.

In OA, miR-133 expression is upregulated in later stages of human joint disease (47) and in synovial fluid and serum signatures associated with OA in human (48) and equine (49) cohorts. Further analysis showed that miR-133 strongly correlated with *HMGB2*, a regulator of chromatin remodeling and fibrosis (48). Mechanistic studies suggest that miR-133 can promote a less viable and less proliferative chondrocyte state. Circ-NCX1 protects chondrocytes from lipopolysaccharide-induced apoptosis by sequestering miR-133a and relieving repression of SIRT1, linking miR-133a to cell survival and autophagy signaling (50). Elevated miR-133a has also been associated with reduced proliferation and a more terminal chondrocyte phenotype (30,50). This is notable because, as mentioned above, regenerative vertebrates reduce miR-133 expression during the proliferative phase of regeneration.

### 5.4. let-7

The let-7 family of microRNAs differs from miR-133 and miR-21 in how it regulates the transition between proliferative and differentiated cell states. Let-7 expression increases during differentiation and senescence but decreases during proliferation and cellular reprogramming (51–54). This dynamic expression pattern curbs cell-cycle progression and is especially relevant during appendage regeneration, where differentiated cells must temporarily regain proliferative and progenitor-like properties. During early regeneration in axolotls, the expression of Lin28—an RNA-binding protein that interacts with let-7 precursors—increases, while let-7 decreases (33,52); let-7 family members are likewise part of the conserved miRNA circuit differentially regulated in regenerating zebrafish, bichir fins and newt tails (32,43). This shift enables the proliferation needed for blastema formation. During this proliferative period, reduced let-7 relieves repression of targets such as IGF1R, allowing cells to respond to growth signals, and acts upstream of the metabolic reprogramming that supports rapid proliferation (52,53). Consequently, prolonged let-7 expression during regeneration may delay blastema formation and inhibit regenerative outgrowth.

In OA, this let-7-mediated cell cycle regulation appears to hinder repair. Instead of transient suppression, some let-7 family members may remain elevated or become induced in diseased joint tissues (55). For example, let-7a-2 increases in the synovial fluid of horses with early OA (55), and let-7b-5p is among the miRNAs enriched in extracellular vesicles released by pro-inflammatory synovial macrophages, a vesicle population that induces cartilage degeneration when transferred to chondrocytes (56). Evidence of intrinsic let-7c suppression in human cartilage suggests that the OA joint retains some capacity for repair (30), although persistent inflammatory injury may prevent this response from progressing to effective regeneration. This capacity varies across joints, with the ankle showing the most permissive state, as with miR-21 described above (30). In the ankle, greater repression of let-7c correlates with higher anabolic turnover, suggesting that reduced let-7 activity may help support the cartilage repair response (30). Supporting studies have demonstrated that suppressing let-7a improves chondrocyte viability and reduces apoptosis, and that inhibiting endogenous let-7a alleviates inflammatory injury by relieving repression of IL6R (57). This effect is congruent across the family: let-7c-5p restrains proinflammatory cytokine production in OA synovial fibroblasts (58), and let-7a-5p delivered to the joint in mesenchymal stromal cell exosomes or platelet-rich plasma promotes chondrocyte proliferation and shifts macrophages toward an anti-inflammatory phenotype (59,60). A key difference between successful regenerative responses in other species and OA is the timing of let-7 suppression. During regeneration, let-7 decreases early, allowing cells to re-enter a proliferative, progenitor-like state, then increases again during differentiation. In OA, persistent let-7 may prevent activated chondrocytes from repairing tissue and producing new matrix. Temporarily suppressing let-7 could overcome this barrier and promote repair. Overall, let-7 links regeneration and OA by regulating whether cells can transition into a repair-supporting state.

## 6. Discussion

### 6.1. MicroRNAs Connect Appendage Regeneration and Osteoarthritis

Comparing appendage regeneration and OA reveals substantial overlap in miRNA programs associated with development, regeneration, and cartilage disease. Rather than acting as isolated regulators of individual processes, miRNAs such as miR-21, miR-133, and let-7, participate in conserved networks regulating cell proliferation, differentiation, metabolism, and tissue remodeling. Their recurrence across developmental and regenerative contexts, as well as in OA, suggests that these pathways may represent shared regulatory machinery that is redeployed during tissue repair and disrupted during disease.

Our identified substantial overlap–of at least 121 miRNAs–is particularly apparent for miR-21 and miR- 133, which are conserved across species and show coordinated changes during regeneration and OA. miR-21 is highly induced during appendage regeneration and is also part of the intrinsic cartilage repair program observed in OA, whereas miR-133 is depleted during early appendage regeneration and increased in late-stage OA. These expression dynamics suggest that human cartilage retains components of regulatory programs active in highly regenerative tissues, but their timing, magnitude, and cellular context may be disrupted. The miRNA let-7 further illustrates how these shared programs can functionally differ. In regeneration, let-7 regulates the transition between differentiated and proliferative states. Its suppression during early regeneration permits cells to enter a repair-competent state, whereas persistent let-7 activity in OA may restrict a reparative response that has already been initiated. Thus, the same regulatory machinery can support regeneration in one context and constrain repair in another, depending on when and where it is active. Our findings indicate that miRNAs are important but underrecognized regulators of both appendage regeneration and degenerative diseases, raising the possibility that similar regulatory mechanisms may extend to other chronic degenerative conditions, including osteoporosis and neurodegenerative diseases.

The extensive overlap of miRNA programs in regenerating and osteoarthritic tissues, as shown in the MiRNA Atlas, offers a catalog of shared regulatory elements. This serves as a framework for identifying regulatory checkpoints that human tissues still possess but may fail to activate, maintain, or resolve properly. Comparative regenerative biology can leverage these conserved miRNA programs not only to identify which factors are present but also to pinpoint which regulatory processes are missing, mistimed, or insufficient in human cartilage, potentially restoring them to improve tissue repair.

### 6.2. The miRNA Atlas as a Resource and its Limitations

The MiRNA Atlas addresses a fundamental gap in comparative skeletal biology by systematically cross-referencing OA and appendage regeneration in these literatures. While the miRNA Atlas identifies shared regulators that would not otherwise emerge from isolated automated literature-mining approaches, it has important limitations that must be acknowledged. First, computational assignment of miRNA regulatory direction relies on natural-language interpretation and may misclassify complex, conditional, or negated findings. Second, the atlas captures reported associations but cannot establish causality or determine whether a miRNA has the same function across contexts. Third, because the atlas is based on published literature, it may be biased toward positive or more readily detected findings and does not capture unpublished or null results. Despite these limitations, the miRNA Atlas provides a systematic and reproducible way to compare miRNAs across OA and appendage regeneration. It enables questions that would otherwise require extensive manual review, including which miRNAs are shared across fields, where they are reported, whether their regulation is consistent, and onto which biological pathways they may converge. In this way, the miRNA Atlas provides a scalable resource for generating and prioritizing hypotheses in regenerative and OA biology.

## 7. Conclusions

Appendage regeneration and OA represent two distinct outcomes of the same injury response. Both begin with cellular reactivation, matrix remodeling, and inflammatory signaling. In regenerating tissue, these signals are coordinated to restore structure and function. In people with clinical manifestations of osteoarthritis, the same regenerative signals persist but fail to overcome injury and restore tissue integrity, resulting instead in progressive degeneration. This literature-derived analysis demonstrates that a substantial portion of the regulatory miRNAs contributing to these divergent outcomes is identical. Therefore, regenerative success and cartilage repair failure may share the same underlying repair machinery but differ in how that machinery is coordinated and whether its activity is sufficient to overcome ongoing injury.

The miRNA Atlas provides a systematic resource for identifying which regulators are shared between these fields and which are unique to each. The three miRNAs examined in greater depth, here, illustrate strong translational potential. Let-7 controls whether cells can transition into a repair-capable state; regenerative appendages open this gate early, while insufficiently repaired osteoarthritic cartilage keeps it closed. MiR-21 and miR-133 regulate proliferation and cell state through pathways shared across regenerative and reparative contexts. By identifying these shared mechanisms, the miRNA Atlas and our accompanying discussion provide concrete, testable targets for potential disease-modifying cartilage therapies. Future work should systematically apply similar analyses to the full set of atlas-identified miRNAs, distinguishing miRNAs that support repair trajectories from those that facilitate disease pathogenesis, and to other regulatory non-coding RNA species (e.g., long non-coding RNAs, piRNAs, circRNAs, etc.). Ultimately, the value of comparative regenerative biology for cartilage therapy lies in learning from species capable of regeneration while identifying the specific barriers that prevent human tissues from achieving the same repair.

## 8. Acknowledgments

This work was supported by the Duke Claude D. Pepper Older Americans Independence Center (NIA P30AG028716). We also thank Davis Zakary for his guidance and support in planning the development and hosting of the application.

## 9. Data Availability Statement

The data used in this study were obtained from publicly available PubMed, Crossref, PMC, and arXiv records. The curated data and associated outputs can be explored through the web application at https://mirnaatlas.shinyapps.io/mirnaatlas/. Source code for the literature-mining application is available from the authors upon reasonable request.

## Notes

https://mirnaatlas.shinyapps.io/mirnaatlas/

### Competing Interest Statement

The authors have declared no competing interest.

## References

1. Haas BJ, Whited JL. Advances in Decoding Axolotl Limb Regeneration. Trends Genet TIG. 2017 Aug;33(8):553–65. doi:10.1016/j.tig.2017.05.006 PubMed PMID: 28648452; PubMed Central PMCID: PMC5534018.

2. McCusker C, Bryant SV, Gardiner DM. The axolotl limb blastema: cellular and molecular mechanisms driving blastema formation and limb regeneration in tetrapods. Regeneration. 2015 May 11;2(2):54–71. doi:10.1002/reg2.32 PubMed PMID: 27499868; PubMed Central PMCID: PMC4895312.

3. Uemoto T, Abe G, Tamura K. Regrowth of zebrafish caudal fin regeneration is determined by the amputated length. Sci Rep. 2020 Jan 20;10:649. doi:10.1038/s41598-020-57533-6 PubMed PMID: 31959817; PubMed Central PMCID: PMC6971026.

4. Wang D, Berg D, Ba H, Sun H, Wang Z, Li C. Deer antler stem cells are a novel type of cells that sustain full regeneration of a mammalian organ—deer antler. Cell Death Dis. 2019 Jun 5;10(6):443. doi:10.1038/s41419-019-1686-y PubMed PMID: 31165741; PubMed Central PMCID: PMC6549167.

5. Yagi S, Hirata M, Miyachi Y, Uemoto S. Liver Regeneration after Hepatectomy and Partial Liver Transplantation. Int J Mol Sci. 2020 Nov 9;21(21):8414. doi:10.3390/ijms21218414 PubMed PMID: 33182515; PubMed Central PMCID: PMC7665117.

6. Michalopoulos GK, Bhushan B. Liver regeneration: biological and pathological mechanisms and implications. Nat Rev Gastroenterol Hepatol. 2021 Jan;18(1):40–55. doi:10.1038/s41575-020-0342-4

7. Shieh S, Cheng T. Regeneration and repair of human digits and limbs: fact and fiction. Regeneration. 2015 Oct 13;2(4):149–68. doi:10.1002/reg2.41 PubMed PMID: 27499873; PubMed Central PMCID: PMC4857729.

8. Mui BWH, Wong JJY, Dumas CE, Wang JH, Bray T, Hirose K, et al. Hyaluronic acid and tissue mechanics orchestrate mammalian digit tip regeneration. Science. 2026 Apr 9;392(6794):eady3136. doi:10.1126/science.ady3136 PubMed PMID: 41955369; PubMed Central PMCID: PMC7619039.

9. Fitzgerald J, Rich C, Burkhardt D, Allen J, Herzka AS, Little CB. Evidence for articular cartilage regeneration in MRL/MpJ mice. Osteoarthritis Cartilage. 2008 Nov 1;16(11):1319–26. doi:10.1016/j.joca.2008.03.014

10. Vovos TJ, Furman BD, Huebner JL, Kimmerling KA, Utturkar GM, Green CL, et al. Initial displacement of the intra-articular surface after articular fracture correlates with PTA in C57BL/6 mice but not “superhealer” MRL/MpJ mice. J Orthop Res Off Publ Orthop Res Soc. 2021 Sep;39(9):1977–87. doi:10.1002/jor.24912 PubMed PMID: 33179316; PubMed Central PMCID: PMC12990664.

11. Lewis JS, Furman BD, Zeitler E, Huebner JL, Kraus VB, Guilak F, et al. Genetic and cellular evidence of decreased inflammation associated with reduced post-traumatic arthritis in MRL/MpJ mice. Arthritis Rheum. 2013 Mar;65(3):660–70. doi:10.1002/art.37796 PubMed PMID: 23203659; PubMed Central PMCID: PMC3721663.

12. Gurtner GC, Werner S, Barrandon Y, Longaker MT. Wound repair and regeneration. Nature. 2008 May;453(7193):314–21. doi:10.1038/nature07039

13. Aztekin C. Appendage regeneration is context dependent at the cellular level. Open Biol. 11(7):210126. doi:10.1098/rsob.210126 PubMed PMID: 34315276; PubMed Central PMCID: PMC8316798.

14. Aztekin C. Tissues and Cell Types of Appendage Regeneration: A Detailed Look at the Wound Epidermis and Its Specialized Forms. Front Physiol. 2021 Nov 23;12:771040. doi:10.3389/fphys.2021.771040 PubMed PMID: 34887777; PubMed Central PMCID: PMC8649801.

15. Vaishya R. The journey of articular cartilage repair. J Clin Orthop Trauma. 2016;7(3):135–6. doi:10.1016/j.jcot.2016.06.001 PubMed PMID: 27489406; PubMed Central PMCID: PMC4949403.

16. Chiang H, Jiang CC. Repair of Articular Cartilage Defects: Review and Perspectives. J Formos Med Assoc. 2009 Feb 1;108(2):87–101. doi:10.1016/S0929-6646(09)60039-5

17. Hoshiyama Y, Otsuki S, Oda S, Kurokawa Y, Nakajima M, Jotoku T, et al. Expression Pattern and Role of Chondrocyte Clusters in Osteoarthritic Human Knee Cartilage. J Orthop Res Off Publ Orthop Res Soc. 2015 Apr;33(4):548–55. doi:10.1002/jor.22782 PubMed PMID: 25691232; PubMed Central PMCID: PMC4454425.

18. Seol D, McCabe DJ, Choe H, Zheng H, Yu Y, Jang K, et al. Chondrogenic Progenitor Cells Respond to Cartilage Injury. Arthritis Rheum. 2012 Nov;64(11):3626–37. doi:10.1002/art.34613 PubMed PMID: 22777600; PubMed Central PMCID: PMC4950521.

19. Tornini VA, Puliafito A, Slota LA, Thompson JD, Nachtrab G, Kaushik AL, et al. Live Monitoring of Blastemal Cell Contributions During Appendage Regeneration. Curr Biol CB. 2016 Nov 21;26(22):2981–91. doi:10.1016/j.cub.2016.08.072 PubMed PMID: 27839971; PubMed Central PMCID: PMC5121098.

20. Currie JD, Kawaguchi A, Traspas RM, Schuez M, Chara O, Tanaka EM. Live Imaging of Axolotl Digit Regeneration Reveals Spatiotemporal Choreography of Diverse Connective Tissue Progenitor Pools. Dev Cell. 2016 Nov 21;39(4):411–23. doi:10.1016/j.devcel.2016.10.013 PubMed PMID: 27840105; PubMed Central PMCID: PMC5127896.

21. Chou CH, Jain V, Gibson J, Attarian DE, Haraden CA, Yohn CB, et al. Synovial cell cross-talk with cartilage plays a major role in the pathogenesis of osteoarthritis. Sci Rep. 2020 Jul 2;10:10868. doi:10.1038/s41598-020-67730-y PubMed PMID: 32616761; PubMed Central PMCID: PMC7331607.

22. Ji Q, Zheng Y, Zhang G, Hu Y, Fan X, Hou Y, et al. Single-cell RNA-seq analysis reveals the progression of human osteoarthritis. Ann Rheum Dis. 2019 Jan;78(1):100–10. doi:10.1136/annrheumdis-2017-212863 PubMed PMID: 30026257; PubMed Central PMCID: PMC6317448.

23. Bartel DP. MicroRNAs: Genomics, Biogenesis, Mechanism, and Function. Cell. 2004 Jan 23;116(2):281–97. doi:10.1016/S0092-8674(04)00045-5

24. Shang R, Lee S, Senavirathne G, Lai EC. microRNAs in action: biogenesis, function and regulation. Nat Rev Genet. 2023 Dec;24(12):816–33. doi:10.1038/s41576-023-00611-y

25. Bartel DP. Metazoan MicroRNAs. Cell. 2018 Mar 22;173(1):20–51. doi:10.1016/j.cell.2018.03.006 PubMed PMID: 29570994; PubMed Central PMCID: PMC6091663.

26. Lee CT, Risom T, Strauss WM. Evolutionary conservation of microRNA regulatory circuits: an examination of microRNA gene complexity and conserved microRNA-target interactions through metazoan phylogeny. DNA Cell Biol. 2007 Apr;26(4):209–18. doi:10.1089/dna.2006.0545 PubMed PMID: 17465887.

27. Mihanfar A, Shakouri SK, Khadem-Ansari MH, Fattahi A, Latifi Z, Nejabati HR, et al. Exosomal miRNAs in osteoarthritis. Mol Biol Rep. 2020 Jun;47(6):4737–48. doi:10.1007/s11033-020-05443-1

28. Szala D, Kopańska M, Trojniak J, Jabłoński J, Hanf-Osetek D, Snela S, et al. The Role of MicroRNAs in the Pathophysiology of Osteoarthritis. Int J Mol Sci. 2024 Jun 8;25(12):6352. doi:10.3390/ijms25126352 PubMed PMID: 38928059; PubMed Central PMCID: PMC11204066.

29. Yu C, Chen WP, Wang XH. MicroRNA in Osteoarthritis. J Int Med Res. 2011 Feb;39(1):1–9. doi:10.1177/147323001103900101

30. Hsueh MF, Önnerfjord P, Kraus VB. Anabolic indices of matrix proteins identify regenerative small RNA intrinsic to human cartilage. Sci Adv. 2025 Jul 11;11(28):eadu8440. doi:10.1126/sciadv.adu8440

31. King BL, Yin VP. Prioritizing Studies on Regeneration in Nontraditional Model Organisms. Regen Med. 2017 Jan;12(1):1–3. doi:10.2217/rme-2016-0159

32. King BL, Yin VP. A Conserved MicroRNA Regulatory Circuit Is Differentially Controlled during Limb/Appendage Regeneration. PloS One. 2016;11(6):e0157106. doi:10.1371/journal.pone.0157106 PubMed PMID: 27355827; PubMed Central PMCID: PMC4927183.

33. Holman EC, Campbell LJ, Hines J, Crews CM. Microarray Analysis of microRNA Expression during Axolotl Limb Regeneration. PLoS ONE. 2012 Sep 13;7(9):e41804. doi:10.1371/journal.pone.0041804 PubMed PMID: 23028429; PubMed Central PMCID: PMC3441534.

34. Zhang Y, Jia J, Yang S, Liu X, Ye S, Tian H. MicroRNA-21 controls the development of osteoarthritis by targeting GDF-5 in chondrocytes. Exp Mol Med. 2014 Feb;46(2):e79. doi:10.1038/emm.2013.152 PubMed PMID: 24577233; PubMed Central PMCID: PMC3944443.

35. Ma S, Zhang A, Li X, Zhang S, Liu S, Zhao H, et al. MiR-21-5p regulates extracellular matrix degradation and angiogenesis in TMJOA by targeting Spry1. Arthritis Res Ther. 2020;22:99. doi:10.1186/s13075-020-2145-y PubMed PMID: 32357909; PubMed Central PMCID: PMC7195789.

36. Konteles V, Papathanasiou I, Tzetis M, Kriebardis A, Tsezou A. Synovial Fibroblast Extracellular Vesicles Induce Inflammation via Delivering miR-21-5p in Osteoarthritis. Cells. 2025 Mar 31;14(7):519. doi:10.3390/cells14070519 PubMed PMID: 40214473; PubMed Central PMCID: PMC11989074.

37. Hoshikawa N, Sakai A, Takai S, Suzuki H. Targeting Extracellular miR-21-TLR7 Signaling Provides Long-Lasting Analgesia in Osteoarthritis. Mol Ther Nucleic Acids. 2019 Nov 20;19:199–207. doi:10.1016/j.omtn.2019.11.011 PubMed PMID: 31841992; PubMed Central PMCID: PMC6920297.

38. Zhu H, Yan X, Zhang M, Ji F, Wang S. miR-21-5p protects IL-1β-induced human chondrocytes from degradation. J Orthop Surg. 2019 May 3;14:118. doi:10.1186/s13018-019-1160-7 PubMed PMID: 31053150; PubMed Central PMCID: PMC6499971.

39. Qin L, Yang J, Su X, Xilan li, Lei Y, Dong L, et al. The miR-21-5p enriched in the apoptotic bodies of M2 macrophage-derived extracellular vesicles alleviates osteoarthritis by changing macrophage phenotype. Genes Dis. 2022 Oct 5;10(3):1114–29. doi:10.1016/j.gendis.2022.09.010 PubMed PMID: 37396516; PubMed Central PMCID: PMC10308169.

40. Chen Y, Yang F, Wang Y, Shi Y, Liu L, Luo W, et al. Mesenchymal stem cell-derived small extracellular vesicles reduced hepatic lipid accumulation in MASLD by suppressing mitochondrial fission. Stem Cell Res Ther. 2025 Mar 5;16:116. doi:10.1186/s13287-025-04228-2 PubMed PMID: 40045380; PubMed Central PMCID: PMC11884000.

41. Yin VP, Thomson JM, Thummel R, Hyde DR, Hammond SM, Poss KD. Fgf-dependent depletion of microRNA-133 promotes appendage regeneration in zebrafish. Genes Dev. 2008 Mar 15;22(6):728–33. doi:10.1101/gad.1641808 PubMed PMID: 18347091; PubMed Central PMCID: PMC2275425.

42. Lepp AC, Carlone RL. RARβ2 expression is induced by the down-regulation of microRNA 133a during caudal spinal cord regeneration in the adult newt. Dev Dyn. 2014 Dec;243(12):1581–90. doi:10.1002/dvdy.24210

43. Lepp AC, Carlone RL. MicroRNA dysregulation in response to RARβ2 inhibition reveals a negative feedback loop between MicroRNAs 1, 133a, and RARβ2 during tail and spinal cord regeneration in the adult newt. Dev Dyn. 2015 Dec;244(12):1519–37. doi:10.1002/dvdy.24342

44. Yu Y, Tang J, Su J, Cui J, Xie X, Chen F. Integrative analysis of microRNAome, transcriptome, and proteome during the limb regeneration of Cynops orientalis.

45. Walker SE, Piazza A, Carlone RL, Spencer GE. MicroRNAs in Tissue Regeneration: Lessons from Animal Models. Int J Mol Sci. 2025 Oct 14;26(20). doi:10.3390/ijms262010043

46. Hutchins ED, Eckalbar WL, Wolter JM, Mangone M, Kusumi K. Differential expression of conserved and novel microRNAs during tail regeneration in the lizard Anolis carolinensis. BMC Genomics. 2016 May 5;17:339. doi:10.1186/s12864-016-2640-3 PubMed PMID: 27150582; PubMed Central PMCID: PMC4858913.

47. Ishida K, Tanishima S, Tanida A, Nagira K, Mihara T, Takeda C, et al. Comprehensive analysis of microRNA expression in lumbar facet joint capsules and synovium of patients with osteoarthritis: Comparison between early-stage and late-stage osteoarthritis samples from a single individual. J Orthop Sci. 2024 Mar 1;29(2):660–7. doi:10.1016/j.jos.2023.01.008

48. Ramos YFM, Coutinho de Almeida R, Lakenberg N, Suchiman E, Mei H, Kloppenburg M, et al. Circulating MicroRNAs Highly Correlate to Expression of Cartilage Genes Potentially Reflecting OA Susceptibility—Towards Identification of Applicable Early OA Biomarkers. Biomolecules. 2021 Sep 13;11(9):1356. doi:10.3390/biom11091356 PubMed PMID: 34572569; PubMed Central PMCID: PMC8468331.

49. Castanheira CIGD, Taylor S, Skiöldebrand E, Rubio-Martinez LM, Hackl M, Clegg PD, et al. Synovial Fluid and Serum MicroRNA Signatures in Equine Osteoarthritis. Int J Mol Sci. 2025 Nov 19;26(22):11190. doi:10.3390/ijms262211190 PubMed PMID: 41303673; PubMed Central PMCID: PMC12652959.

50. Liu K, Fan X, Zhang L, Yang Y, Zhou X. Circ-NCX1 inhibits LPS-induced chondrocyte apoptosis by regulating the miR-133a/SIRT1 axis. Kaohsiung J Med Sci. 2022 Jul 27;38(10):992–1000. doi:10.1002/kjm2.12564 PubMed PMID: 35894157; PubMed Central PMCID: PMC11896150.

51. Wang Y, Zhao J, Chen S, Li D, Yang J, Zhao X, et al. Let-7 as a Promising Target in Aging and Aging-Related Diseases: A Promise or a Pledge. Biomolecules. 2022 Aug 2;12(8):1070. doi:10.3390/biom12081070 PubMed PMID: 36008964; PubMed Central PMCID: PMC9406090.

52. Varela-Rodríguez H, Abella-Quintana DG, Espinal-Centeno A, Varela-Rodríguez L, Gomez-Zepeda D, Caballero-Pérez J, et al. Functional Characterization of the Lin28/let-7 Circuit During Forelimb Regeneration in Ambystoma mexicanum and Its Influence on Metabolic Reprogramming. Front Cell Dev Biol. 2020 Nov 19;8:562940. doi:10.3389/fcell.2020.562940 PubMed PMID: 33330447; PubMed Central PMCID: PMC7710800.

53. Hu W, Li T, Hu R, Wu L, Li M, Meng X. MicroRNA let-7a and let-7f as novel regulatory factors of the sika deer (Cervus nippon) IGF-1R gene. Growth Factors. 2014 Feb 1;32(1):27–33. doi:10.3109/08977194.2013.860453

54. Johnson CD, Esquela-Kerscher A, Stefani G, Byrom M, Kelnar K, Ovcharenko D, et al. The let-7 MicroRNA Represses Cell Proliferation Pathways in Human Cells. Cancer Res. 2007 Aug 15;67(16):7713–22. doi:10.1158/0008-5472.CAN-07-1083

55. Castanheira C, Balaskas P, Falls C, Ashraf-Kharaz Y, Clegg P, Burke K, et al. Equine synovial fluid small non-coding RNA signatures in early osteoarthritis. BMC Vet Res. 2021 Jan 9;17:26. doi:10.1186/s12917-020-02707-7 PubMed PMID: 33422071; PubMed Central PMCID: PMC7796526.

56. Zhao S, Wang J, Xue M, Wu B, Sheng L, Wen Y, et al. Synovial inflammatory macrophage-derived extracellular vesicles exacerbate cartilage lesions with a FMRP-selectively sorted manner in osteoarthritis. Bone Res. 2026 Feb 17;14:26. doi:10.1038/s41413-025-00502-4 PubMed PMID: 41702879; PubMed Central PMCID: PMC12913794.

57. Sui C, Zhang L, Hu Y. MicroRNA-let-7a inhibition inhibits LPS-induced inflammatory injury of chondrocytes by targeting IL6R. Mol Med Rep. 2019 Sep;20(3):2633–40. doi:10.3892/mmr.2019.10493 PubMed PMID: 31322277; PubMed Central PMCID: PMC6691277.

58. Law YY, Lee WF, Hsu CJ, Lin YY, Tsai CH, Huang CC, et al. miR-let-7c-5p and miR-149-5p inhibit proinflammatory cytokine production in osteoarthritis and rheumatoid arthritis synovial fibroblasts. Aging. 2021 Jul 1;13(13):17227–36. doi:10.18632/aging.203201 PubMed PMID: 34198264; PubMed Central PMCID: PMC8312412.

59. Shao L, Ding L, Li W, Zhang C, Xia Y, Zeng M, et al. Let-7a-5p derived from parathyroid hormone (1–34)-preconditioned BMSCs exosomes delays the progression of osteoarthritis by promoting chondrocyte proliferation and migration. Stem Cell Res Ther. 2025 Jun 9;16:299. doi:10.1186/s13287-025-04416-0 PubMed PMID: 40490830; PubMed Central PMCID: PMC12150519.

60. Li Q, Wang M, Wang D, Nie Y, Sun X, Na L, et al. Platelet-rich plasma-derived microRNA let-7a-5p alleviates knee osteoarthritis by regulating macrophage polarization and improving inflammatory microenvironment. Front Immunol. 17:1756467. doi:10.3389/fimmu.2026.1756467 PubMed PMID: 41777883; PubMed Central PMCID: PMC12950549.

